# Humanization of *Drosophila* Gαo to model *GNAO1* paediatric encephalopathies

**DOI:** 10.1101/2020.08.14.251173

**Authors:** Mikhail Savitsky, Gonzalo P. Solis, Vladimir L. Katanaev

**Affiliations:** Translational Research Center in Oncohaematology, Department of Cell Physiology and Metabolism, Faculty of Medicine, University of Geneva, Geneva, Switzerland; School of Biomedicine, Far Eastern Federal University, Vladivostok, Russia

## Abstract

Several hundred genes have been identified to contribute to epilepsy – the disease affecting 65 million people worldwide. One of these genes is *GNAO1* encoding Gαo, the major neuronal α-subunit of heterotrimeric G proteins. An avalanche of dominant *de novo* mutations in *GNAO1* have been recently described in paediatric epileptic patients, suffering in addition to epilepsy from motor dysfunction and developmental delay. Although occurring in amino acids conserved from humans to *Drosophila*, these mutations and their functional consequences have only poorly been analysed at the biochemical or neuronal levels. Adequate animal models to study molecular aetiology of *GNAO1* encephalopathies have also so far been lacking. As the first step towards modelling the disease in *Drosophila*, we here describe humanization of the *Gαo* locus in the fruit fly. A two-step CRISPR/Cas9-mediated replacement was conducted, first substituting the coding exons 2-3 of *Gαo* with respective human *GNAO1* sequences. At the next step, the remaining exons 4-7 were similarly replaced, keeping intact the gene *Cyp49a1* embedded in-between, as well as the non-coding exon 1 and the surrounding regulatory sequences. The resulting flies, homozygous for the humanized *GNAO1* loci, are viable and fertile without any visible phenotypes; their body weight and longevity are also normal. Human Gαo-specific antibodies confirm the endogenous-level expression of the humanized Gαo, which fully replaces the *Drosophila* functions. The genetic model we established will make it easy to incorporate encephalopathic *GNAO1* mutations and will permit intensive investigations into the molecular aetiology of the human disease through the powerful toolkit of *Drosophila* genetics.

## Introduction

Epilepsy is a chronic disease of multigenic origin, characterized by appearance, mostly in an unpredicted manner, of seizures, and sometimes complicated by other neurological or neurodevelopmental deficits (Staley, 2015). Seizures result from an imbalance between inhibitory and excitatory conductances in the brain and can be induced by acute toxic or traumatic impacts. The causes of the episodic shifts in the balance of inhibition and excitation seen in the chronic epilepsy remain largely unexplained. With 65 millions of people worldwide currently suffering of epilepsy and the inadequacy of the current pharmacological approaches to certain subtypes of the disease, the need to advance and complete our understanding of the aetiology of this disease is clear, as is the urgency to develop novel medical treatments (Krook-Magnuson and Soltesz, 2015).

Several hundred genes linked to epilepsy have been identified (Ran et al., 2015), and the estimate tells that this number will increase by folds, matching the number of genes linked to cancer and exceeding the diversity of the cancer-related genes by biological functions (Grone and Baraban, 2015; Noebels, 2015). Advances have been made in the understanding of the epileptic molecular and cellular mechanisms and pathways (involving ion channels and their regulators, but also neuronal migration, synaptic plasticity, and neurite outgrowth, among other things (McTague et al., 2016; Noebels, 2015; Staley, 2015)). However, many of the epileptic mutations remain enigmatic – both in the sense of how they provoke the disease, as well as in the sense of the broader molecular pathway, hijacked in epilepsy, of which they are part. One of the proteins who has recently emerged as an important player in epilepsy is Gαo – the major neuronal α-subunit of heterotrimeric G proteins, encoded by the gene *GNAO1*. Starting from 2013 (Nakamura et al., 2013), whole-exome sequencing in a subset of epileptic patients has resulted in an avalanche of described mutations in *GNAO1* (see (Schirinzi et al., 2019) for the most up to date review). This subset represents paediatric patients with the early onset epileptic encephalopathy and Ohtahara syndrome, suffering additionally from movement disorders and developmental delays. These patients are resistant to conventional antiepileptic pharmacological treatments and may die early in life. A number of sites in Gαo have been found mutated in these epileptic patients, all affecting amino acids conserved among Gαo and its orthologues all the way down to fruit flies and nematodes (Fig. 1A), highlighting the importance of the affected amino acids for the Gαo function. *GNAO1* mutations in epilepsy are heterozygous suggesting their dominant nature. Interestingly, the relative degree of manifestation of different deficits (seizures, developmental delay, movement disorders) may be different depending on the exact amino acid mutation in Gαo (Menke et al., 2016; Schorling et al., 2017; Talvik et al., 2015). Recently, a point mutation in Gαo was identified to cause a severe childhood speech disorder (Hildebrand et al., 2020).

**Figure 1.**
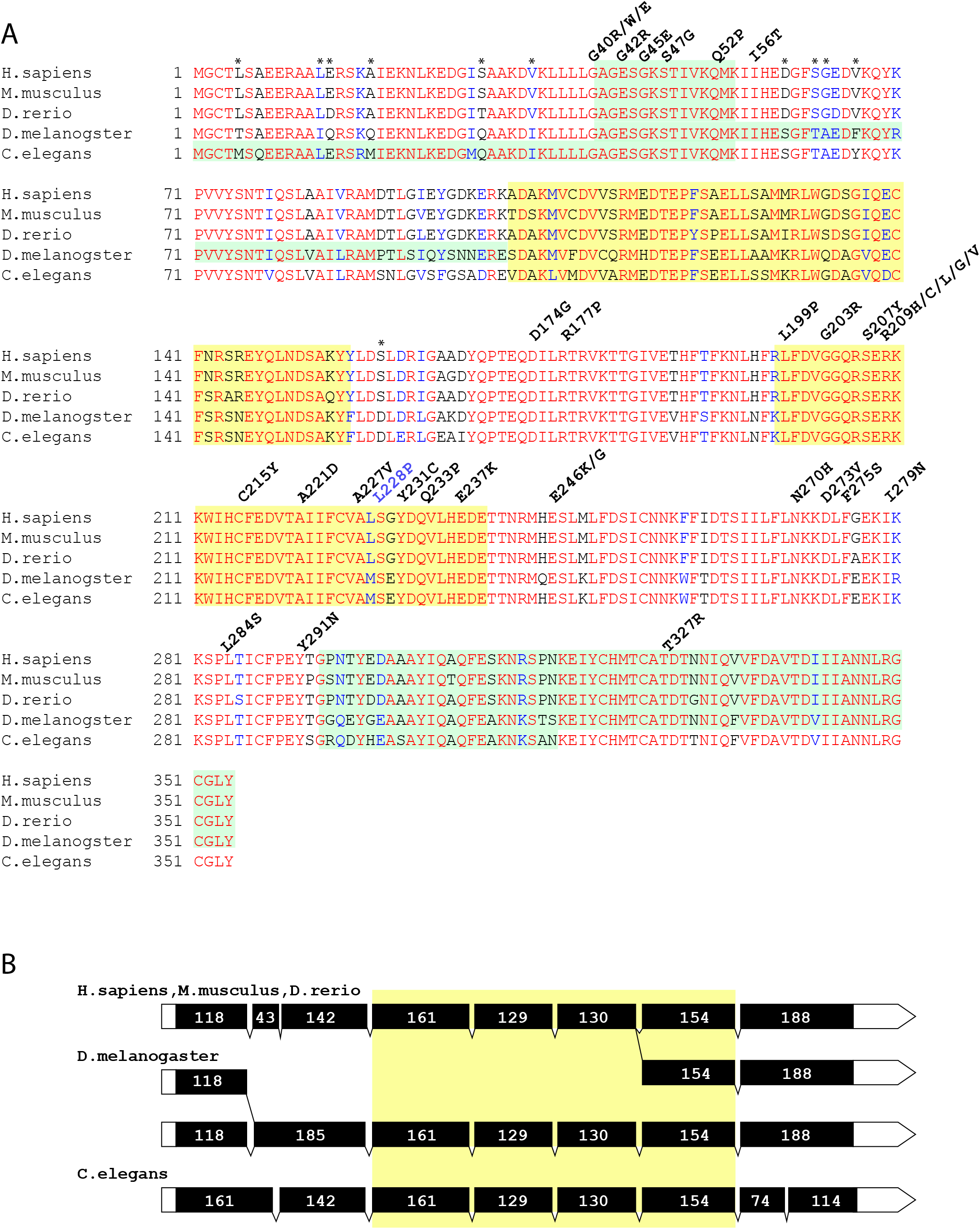
A. Alignment of Gαo homologs from three vertebrate and two invertebrate organisms. Pale green and yellow colours on the background of the alignment mark exon boundaries. Above the alignment, the known mutations presented in human genomes and associated with human neurological phenotypes are designated. Mutated amino acids belong to conservative parts of proteins (letter blocks of red). The mismatching amino acids which were not replaced, during editing dGαo are designated with asterisks. For *Drosophila*, the dGαo-B isoform is shown. B. Schematic exon-intron structures of Gαo homologs. Black boxes mark the coding parts of the exons and white boxes indicate the noncoding parts. The drawing of the exons is presented in scale. Digits inside exons mark their precise length in base pairs. The yellow box marks identical exons in compared species.

Heterotrimeric G proteins are key signalling molecules best recognized as the immediate transducers of GPCRs (G protein coupled receptors). Coupled to GPCRs, heterotrimeric G proteins transduce signals from a large variety of extracellular cues, from quanta of light, ions, small organic molecules, to large macromolecules (Fredriksson and Schioth, 2005). A heterotrimeric G protein complex consists of three subunits, α, β, and γ, of which the α-subunit is responsible for binding to guanine nucleotides as well as to the cognate GPCR. Sixteen vertebrate Gα subunits have been identified and classified into four major families based on sequence homology: Gαi/o, Gαs, Gαq and Gα12 (Milligan and Kostenis, 2006). Gαo belongs to the first family and transduces the signal of a group of rhodopsin-like GPCRs including the opioid, α2-adrenergic, D2 dopaminergic, M2 muscarinic, and somatostatin receptors (Wettschureck and Offermanns, 2005). Gαo was among the first α-subunits discovered (Sternweis and Robishaw, 1984) and found to be the major Gα subunit of the central nervous system (CNS) across the animal kingdom (Strathmann and Simon, 1990; Wolfgang et al., 1990), controlling both development and adult physiology of the brain (Greif et al., 2000; Jiang et al., 1998) and the main olfactory system (Choi et al., 2016; Oboti et al., 2014; Tanaka et al., 1999). Gαo knockout (KO) mice showed a strong developmental delay during the first 3 weeks after birth, and a short half-life of only 7 weeks on average (Jiang et al., 1998). Gαo KO mice also presented multiple neurological abnormalities such as hyperalgesia, hyperactivity, generalized tremor with occasional seizures, and a severe impairment of motor control (Greif et al., 2000; Jiang et al., 1998). At the cellular levels, dorsal root ganglion cells derived from Gαo KO mice presented a reduced inhibition of Ca^2+^ channel currents by the agonist-induced activation of opioid receptors (Jiang et al., 1998). Gαo has also been implicated in the regulation of Ca^2+^ and K^+^ channels in sensory and hippocampal neurons (Campbell et al., 1993; Greif et al., 2000). In addition to neurons, Gαo is expressed in the heart (Strathmann et al., 1990) where it is necessary for proper development and functioning of the organ (Fremion et al., 1999; Valenzuela et al., 1997). Gαo has additional developmental functions such as cell fate determination and polarization, in part via the Wnt-Frizzled pathway (Egger-Adam and Katanaev, 2008; Koval et al., 2011), as well as pathological implications e.g. in cancer (Entschladen et al., 2011).

Gαo expression was found to strongly increase during early neonatal rat brain development (Milligan et al., 1987) where it is enriched in growth cones (Strittmatter et al., 1990). In the rat pheochromocytome cell line PC12, Gαo expression rises during the process of neurite outgrowth induced by NGF (Andreopoulos et al., 1995; Asano et al., 1989; Zubiaur and Neer, 1993), and overexpression of a Gαo active mutant potentiates the effects of NGF in neurite length (Strittmatter et al., 1994). Similar to its mammalian counterpart, insect Gαo is strongly expressed in the CNS of the adult *Drosophila* (Wolfgang et al., 1990). Gαo transcript and protein were present at all stages of embryonic development with a marked increase during the period of active axonogenesis (Guillen et al., 1991; Wolfgang et al., 1991). These early data suggested that Gαo may be involved in neuronal differentiation, a role supported by later studies showing reduced neurogenesis and increased cell death in the olfactory bulb of Gαo KO mice (Choi et al., 2016) and defects in guidance and axonal growth of motoneurons in Gαo mutant fruit flies (Fremion et al., 1999). Additionally, differentiation of mouse embryonic stem cells into dopaminergic neurons pointed to Gαo as one of the genes specifically upregulated during neurogenesis (Lee et al., 2006).

Driven by the pivotal roles of Gαo and by the fact that the list of its molecular targets was remarkably short (Jiang and Bajpayee, 2009) we have performed massive whole genome/proteome screenings to identify novel Gαo interaction partners. This analysis identified >250 proteins as novel candidate Gαo partners. “Cherry-picking” of individual proteins from this list resulted in detailed descriptions of novel mechanisms of Gαo-controlled regulation of Wnt/Frizzled signalling, neuromuscular junction formation, planar cell polarity, asymmetric cell divisions, *etc*. (Egger-Adam and Katanaev, 2010; Kopein and Katanaev, 2009; Lin and Katanaev, 2013; Lin et al., 2014; Luchtenborg et al., 2014; Purvanov et al., 2010). As opposed to such characterizations of selected individual Gαo partners, we next aimed at identifying functional modules within the Gαo interactome. Several functional modules (such as cytoskeleton organization, cell division, cell adhesion, *etc*.) were identified within the Gαo interactome; subsequent work identified Gαo as a master regulator of vesicular trafficking (Solis et al., 2017). Remarkably, across different cell types (neuronal, mesenchymal, epithelial) and species (insects and mammals), Gαo is found to dually localize to PM (plasma membrane) and Golgi. This dual localization is found to play a coordinated role in formation of cellular protrusions (such as neurites in neuronal cells), so that the PM pool of Gαo indices the initiation of the protrusion, while the Golgi pool ensures material delivery to it, permitting its elongation and stabilization. In the nervous system, these novel Gαo activities are necessary not only for neuritogenesis, but also for synaptogenesis (Solis et al., 2017). As improper synaptic plasticity and process outgrowth are both implicated in the aetiology of epilepsy (Staley, 2015), the possible dominant effects of the epileptic Gαo mutations on these novel Golgi-emanating mechanisms discovered by us need to be investigated in detail, as are the molecular events happening at PM. Further, with several established epileptic mutations affecting proteins working in Golgi-mediated trafficking (http://61.152.91.49/EpilepsyGene/), and likely more to emerge, such investigation could be the entry point into identification of a novel epileptic pathway, involving Gαo-mediated trafficking routes.

Of the point mutations identified within the coding region of *GNAO1* in paediatric encephalopathy patients, all correspond to highly conserved residues (Fig. 1A), indicating their involvement in basic Gαo functions. However, molecular characterization of these mutants has so far been very limited and provide to a certain degree contradicting findings on mutant protein expression in heterologous cells, the effects of the mutants on basal and norepinephrine-induced N-type calcium channels, and on the forskolin-stimulated cAMP production (Feng et al., 2017; Nakamura et al., 2013). Thus, the molecular mechanism(s) of Gαo mutants in the development of the severe neurological disorders awaits the much-needed detailed clarification to reveal potential therapeutic targeting approaches.

Animal models have the instrumental role in deciphering the disease mechanisms and in identifying / validating the treatment routes. *Drosophila* has been used to model a variety of human maladies, especially neurological ones (Bellen et al., 2019; Bolus et al., 2020; Katanaev et al., 2020; Katanaev and Kryuchkov, 2011; Luchtenborg and Katanaev, 2014). As the first step towards modelling *GNAO1* paediatric encephalopathies in the fruit fly, we here describe humanization of the *Drosophila* Gαo locus, finding the human protein fully replaces the insect one’s functions without any aberrations.

## Results and Discussion

Comparative analysis of Gαo homologs reveals a high degree of similarity in the amino acid sequences of proteins as well as in the exon-intron structures of the genes encoding them. Even evolutionary distant organisms keep the same nucleotide length of the coding sequences in alternatively spliced transcripts and have the resultingly fixed protein length: 354aa. However, lengths of the genomic loci vary in a wide range from species to species. In humans, the respective locus extends for 165kb, in *Drosophila* – for 28kb, and in *C. elegans* – for 4.5kb (see the gene links ncbi.nlm.nih.gov/gene/?term=2775, flybase.org/reports/FBgn0001122 (also see Fig.2A), and wormbase.org/species/c_elegans/gene/WBGene00001648#0-9f-10, respectively, Supplementary Fig. 1). Interestingly, all of them have very similar sets of coding exons (Fig. 1B). For example, 191 amino acids between 101aa and 292aa are coded by 4 exons with the lengths 161, 129, 130 and 154bp in both invertebrates and humans. The percent of similarity between human and *C. elegans* Gαo proteins is 86.7%, with 82.5% identity. The *Drosophila* protein is slightly more similar to the human one: 86.5% similarity and 83.9% identity, with only 57 mismatching amino acids out of 354. The alignment of 5 protein sequences (two highly similar Gαo isoforms are encoded from the same gene both in humans and in *Drosophila*) reveals the presence of conservative and independently evolving blocks, which have not changed through hundreds of millions of years (Fig. 1B).

With this level of conservation, it is perhaps not surprising that of the *de novo* point mutations so far identified in patients suffering from *GNAO1* encephalopathies (altogether 34 different point mutations occurring in 27 sites), all but one fall into the amino acids identical between human, *Drosophila* and nematode Gαo sequences (Fig. 1A). This fact suggests that these amino acids play instrumental functions for the activity of Gαo, and also provides the ground to attempt modelling of *GNAO1* encephalopathy in *Drosophila*. For the sake of terminology, we will be referring to the *Drosophila* and human genes as *Gαo* and *GNAO1*, and to the proteins they encode – as dGαo and hGαo, respectively.

*Drosophila Gαo* produces two variants of the protein through alternative splicing of first coding sequence-containing exons. The six downstream exons are common for both transcripts (Fig. 1B). These splice variants differ only in 7 amino acids, making them 98% identical to each other. In pairwise comparison with the two isoforms, hGαo is closer to the dGαo-B isoform (Supplementary Fig. 2). Both variants of dGαo are expressed at similar levels during the life cycle, with one minor exception: one of the isoforms starts to be transcribed in very early embryos and the other one – 10 hours later after the beginning of embryogenesis (Wolfgang et al., 1991). The two alternatively spliced first coding exons are distanced from each other by 6kb. The six downstream common exons are grouped into two clusters with 2 and 4 exons, which are separated by a long intron containing embedded gene *Cyp49a1*, oriented in the direction opposite to *Gαo* (Fig. 2).

**Figure 2.**
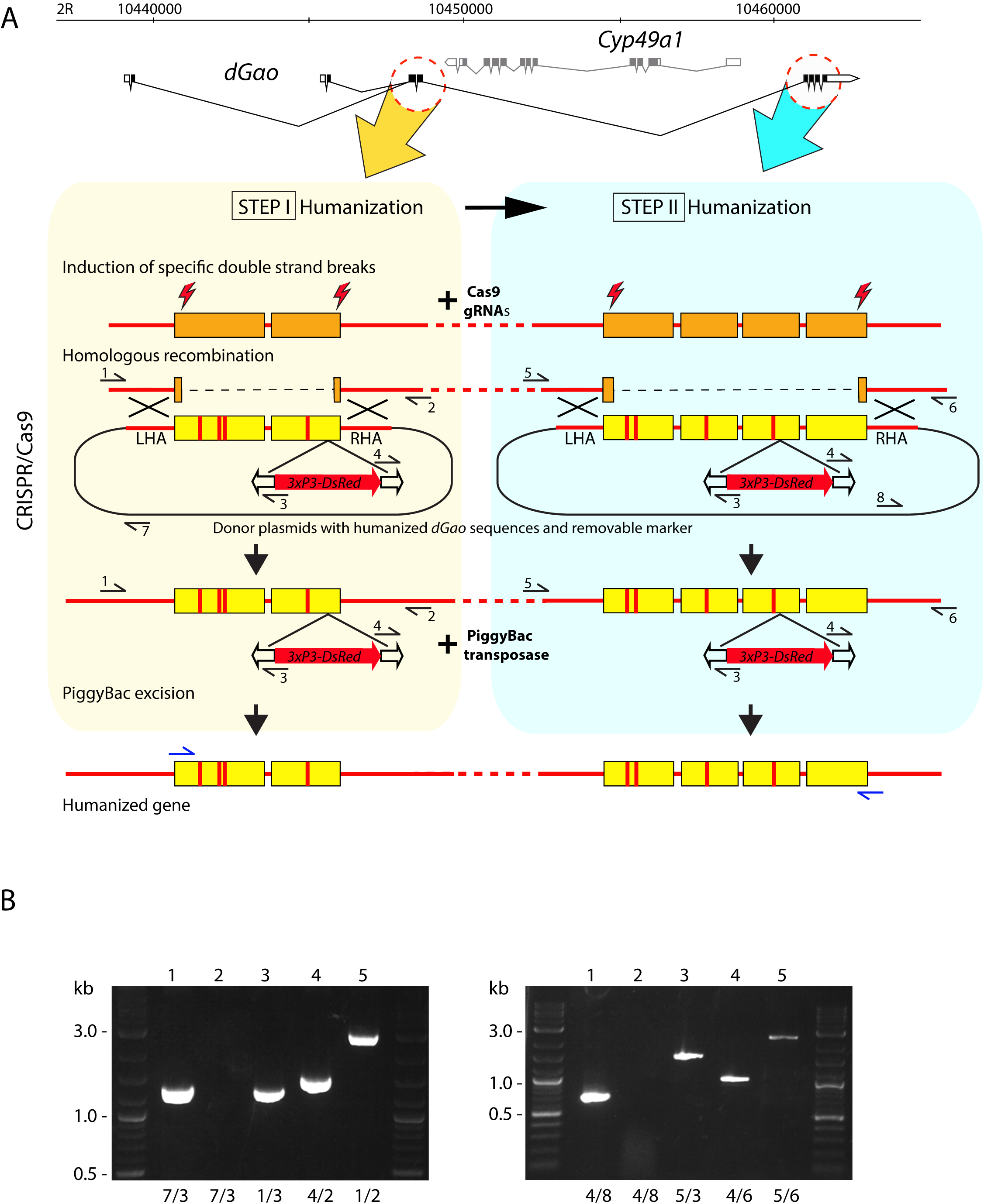
A. The two-step strategy for genomic humanization of dGαo. Primers for validations of correct donor sequences integration in the genome are designated with black horizontal arrows and marked with digits. Primers used in RT-PCR are designated with blue horizontal arrows. LHA and RHA: left and right homologous arms in the donor plasmids, required for homologous recombination with genomic sequences. Exons with vertical red lines schematically designate humanized sequences with mismatches. B. Agarose gels with PCR products obtained with the primers as numbered below each lane (see Supplementary Table 1 for sequences). Lanes 1: control PCR from donor plasmids pLdhGao23R (left panel) and pLdhGao47R (right panel). The absence of a PCR product (lanes 2) performed with the same primers from genomic DNA of transgenic flies indicates the correct integration of the donor sequences without the plasmid backbone. Lanes 3 and 4: verification of the entire integration of the donor sequence in the genome before excision of the 3xP3-dsRed marker. Lanes 5: PCR through the integrated donor sequence after excision of the 3xP3-dsRed marker from genomic DNA of the homozygous stocks *Gαo[h23ex4aa-1]* (left panel) and *Gαo[humanized-1]* (right panel).

The locus of human *GNAO1* contains a duplication of the two last exons, which are spliced alternatively to produce two protein isoforms (hGαoA and hGαoB) with amino acids variations in the C-terminus. In pairwise comparison with the two human isoforms, dGαo is closer to the hGαoB isoform (Supplementary Fig. 2). Unlike the two *Drosophila* isoforms, the two human splice variants of *GNAO1* are expressed differently, with hGαoA being the major version expressed (Consortium, 2013). All known mutations (from E246K to Y291N) (Fig. 1A) in the alternatively spliced exons 7 and 8 belong to hGαoA. Thus, we concentrated on this variant in our replacement strategy.

In order to preserve the endogenous structure of the *Gαo* locus with its potential regulatory elements such as enhancers, promoters, insulators *etc*., we developed the strategy of sequential *Gαo* editing during the humanization process (Fig. 2A). In order to ensure the non-damaged *Gαo* transcription start sites, correct splicing with production of the two *Gαo* transcripts with endogenous and undamaged 5’UTR sequencing, this strategy involved keeping untouched the non-coding exon together with the first coding exon, which is subject to the alternative splicing. The sequences of the core of the gene were humanized in two steps. In the first step, we humanized the exons 2 and 3; in the second – exons 4 to 7 (Fig. 2A). Both rounds of transgenesis were performed with the CRISPR-Cas9 technology (see Methods for details). As a template for the homologous recombination, we used donor plasmids containing *Drosophila* DNA sequences, whereas the codons different between *Gαo* and *GNAO1* contained the minimally needed nucleotide substitutions ensuring encoding of the human amino acids. Additionally, the donor plasmid contained synonymous nucleotide substitutions in the sequences corresponding to the Cas9/gRNA complex targets preventing their recognition and destruction by the complex.

For the gene editing, we used the transgenic fly strain expressing Cas9 in the germline. In order to replace the first cluster of exons, we injected embryos with the mixture of 4 plasmids: the donor plasmid for homologous recombination and 3 plasmids producing gRNAs for the induction of breaks in the target genome locus (see Methods). Besides the modified *Gαo* exons and the homologous arms (LHA and RHA), the donor plasmids contained a 3xP3-DsRed marker cassette flanked by PiggyBac transposon ends inserted in the end of the 3^rd^ exon. Transgenic flies were selected by red fluorescence in eyes provided by expression of DsRed under the eye-specific 3xP3 promoter (Berghammer et al., 1999). Established transgenic fly strains were verified by PCR with primers annealing to the neighbouring genomic sequences outside of the homologous arms and inside the PBac transposon. Correct sizes of PCR products indicated proper integration of the donor plasmid (Fig. 2B). Fly strains with integration of the whole donor plasmid were excluded from the analyses by using primers annealing at the body of plasmid. About one third of the transgenic lines contained the plasmid integrated via the rolling circle replication mechanism and thus had to be discarded.

Three independent transgenic lines with the full set of substitutions, and two independent transgenic lines lacking 4aa substitutions in the 2^nd^ exon (see below for the explanation on the 4aa) were identified after the sequencing analysis of the PCR products and selected for further propagation. These fly lines were crossed with the strain encoding the PBac transposase. The excision of the marker gene was identified in the second generation by loss of red fluorescence in eyes. Since mobilization of PBac-based transposons is characterized by precise excisions without indels in the insertion site, the integrity of the 3^rd^ exon was restored and, consequently, this partially humanized gene started producing proper transcripts. Finally, three independent strains with the full set of substitutions in the first exon cluster, along with the two independent strains without 4aa substitutions in the 2^nd^ exon, were established. These alleles were named *Gαo[h23ex-1], Gαo[h23ex-2], Gαo[h23ex-3]*, and *Gαo[h23ex4aa-1], Gαo[h23ex4aa-2]*, respectively. Both *Gαo[h23ex4aa]* lines demonstrated clear viability and fertility in homozygosity. In a sharp and curious contrast, all the three *Gαo[h23ex]* lines were homozygous lethal.

After removing the marker gene both allele types were confirmed by PCR and sequencing. Then we extracted total RNA from adult flies, and *Gαo* transcripts were examined through RT-PCR using the primers for wild-type and mutant alleles. PCR products amplifying the 958bp region between exons 2 and 7 exons were verified by sequencing, revealing that all the mutant transcripts were spliced correctly, and, as expected, that both the wild type and the mutant transcripts were present in the heterozygous *Gαo[h23ex]* stocks. Homozygous *Gαo[h23ex4aa]* stocks produced mutant transcripts only. Due to the unexplained lethality of the *Gαo[h23ex]* flies, the second round of humanization was performed on the *Gαo[h23ex4aa-1]* strain.

The second cluster was edited in the same manner as the first one: the *Gαo[h23ex4aa-1]* stock combined with the Cas9-expressing transgene was used for the germline transformation (see Methods). Transgenic flies were selected by the fluorescent marker, and then verified by PCR (Fig. 2B) followed by sequencing both before and after excision of the marker; ultimately, the resultant transcript sequences were also verified. Two independent fly lines obtained through the two-step humanization process were established. In these lines, 26 out of the 27 amino acids different between the human and *Drosophila* Gαo were replaced (except of F156Y, apparently not substituted as a result of gRNA imperfection, adding this non-replaced amino acid to the four not replaced in the first round of humanization). These alleles were named *Gαo[humanized-1]* and *Gαo[humanized-2]*. Both lines were homozygous viable and fertile.

As a result of this two-step CRISPR/Cas9-mediated humanization, we succeeded to replace, in the endogenous *Gαo* locus, 49 amino acids of dGαo with the human Gαo sequences. The resultant identity between hGαo and the humanized *Gαo* isoform dGαo-A reached 96.8%, and the humanized isoform dGαo-B – 97.6%. These values are similar to those between the mouse and human Gαo sequences and are much higher than within any vertebrate-invertebrate Gαo pair.

In addition to validation of the humanized transcript expression (Fig. 2B), we confirmed hGαo expression in *Drosophila* central nervous system by immunostaining. Brains together with ventral nerve cords from third instar larvae were probed with antibodies against dGαo (polyclonal) and hGαo (monoclonal, see Methods). We found that the polyclonal antibodies expectedly recognized both *Drosophila* and human proteins in our samples, while the monoclonal antibodies had affinity exclusively to the human Gαo (Fig. 3A). The pattern of immunostaining with different antibodies was identical for both genotypes. These findings support our conclusion that hGαo is expressed in a proper way under the endogenous *Gαo* gene regulatory elements. This expression of hGαo was sufficient to fully recapitulate the endogenous functions of dGαo, as judged by the fertility of the humanized lines and their full viability, measured e.g. by their normal body weight (Fig. 3B) and life span (Fig. 3C).

**Figure 3.**
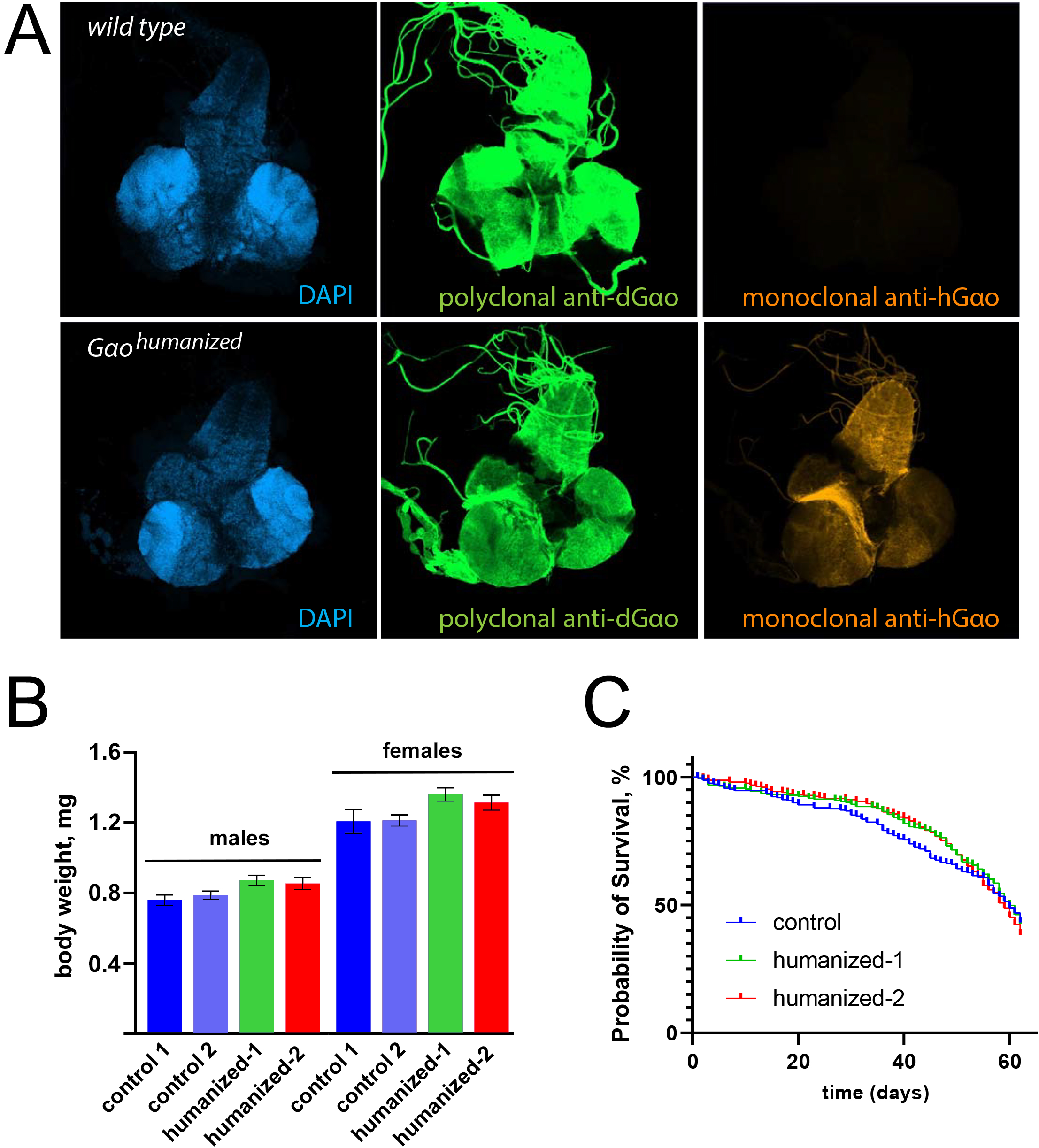
A. Expression of humanized and wild-type Gαo in brains and ventral nerve cords of 3rd instar larvae. The upper panel represents a sample from wild-type larvae and the bottom panel – from *dGao[humanized]*. The immunostaining demonstrates specific recognition of the humanized protein (bottom right) with the monoclonal antibody recognizing hGαo; the polyclonal anti-dGαo antibodies, in contrast, recognize both dGαo and hGαo (the latter with a lower affinity as seen in the middle panels, which were recorded with identical confocal settings). B. Body weight of the humanized flies is normal. The body weight in two control lines (the line used for initial injection is control 1 and an independent line is control 2) and the two homozygous humanized lines generated in our study was determined through measuring the weight of groups of fifty adult (2-3 days after hatching) male or female flies kept under non-crowding conditions; the individual fly body weight was then recalculated. Data are shown as mean ± sd, n= 11 to 13 groups. C. Life span of 250 adult flies (125 males and 125 females; control animals were of the same genotype as ‘control 1’ in (B)) was determined for over 60 days.

Replacement of a model organism’s protein-coding sequence with its human ortholog, preserving the endogenous gene expression regulators, represents an important tool to investigate the evolutionary and functional conservation of the gene of interest; this approach is particularly important for pathology-related genes (Takano-Shimizu-Kouno and Ohsako, 2018). A systematic investigation into the efficiency of gene humanization has been performed in yeast (Kachroo et al., 2015). Despite several successful investigations in *Drosophila* (Chang and Morton, 2017; Wangler et al., 2017), the fruit fly has not been systematically studied in this regard. Our study describes the first ever replacement of an endogenous *Drosophila* G protein with the orthologous human sequence. We find it remarkable that humanized Gαo fully recapitulates the numerous *Drosophila* protein functions, resulting in phenotype-less, viable and fertile flies, having the normal life cycle and longevity. The resultant *Drosophila* strain humanized for the *Gαo* gene can now be used to incorporate human mutations found in *GNAO1* encephalopathy patients. The full power of *Drosophila* genetics and drug discovery tools available in this model organism (Sonoshita and Cagan, 2017; Yamaguchi, 2018) will then become accessible to uncover the molecular mechanisms of the aetiology of *GNAO1* encephalopathy and to provide potential therapeutic leads to treat it. The progressive manner of this devastating disease affecting infants and the current lack of efficient treatment makes this approach highly desired.

## Materials and Methods

### Plasmids

#### Donor plasmid pLdhGao23R for the first round of humanization dGαo

DNA fragment of humanized *Drosophila Gαo* containing exons 2 and 3 with the intron between them was ordered in gene synthesis company (Synbio Technologies) as the plasmid pUC57-dhGao23. The synthesized fragment contained nucleotide substitutions in the *Drosophila* sequence minimally needed to encode human amino acids. Additionally, synonymous substitutions were introduced in the target sites for the gRNA-Cas9 complex in order to avoid double-strand breaks. The fragment was further designed to contain AarI and EcoRV restriction sites adjoining the exon 2, and an additional AarI site directly after the exon 3.

The left homologous arm (LHA) was PCR-amplified with the LHAdGao23fw and LHAdGao23rev primers from genomic DNA using Phusion High-Fidelity DNA Polymerase (New England Biolabs), producing 1060bp PCR product, which was further cloned into the pUC57-dhGao23 plasmid by the EcoRV site (producing the construct pLdhGao23). The right homologous arm (RHA) was PCR-amplified with the RHAdGao23fw and RHAdGao23rev primers, and the resulting 1070bp PCR product was cloned into the plasmid pHD-ScarlessDsRed (DGRC,1364) into the SapI (producing the construct pScarless-dhGaoR). Both cloning steps were performed with the NEBuilder HiFi DNA Assembly Cloning Kit (New England Biolabs). The AarI-AarI fragment from pLdhGao23 was cloned into the AarI site of pScarless-dhGaoR. The resultant donor plasmid pLdhGao23R contains humanized exons 2 and 3 of *Gαo* flanked with 1000bp-long LHA and RHA. The end of exon 3 of *Gαo* in the plasmid is modified by insertion of the piggyBac transposon between duplicated TTAA sequence. Precise excision of piggyBac restores correct exon sequence with unique TTAA.

#### Donor plasmid for the second round of humanization dGαo

DNA fragment of humanized Drosophila *Gαo* containing exons 4 to 7 with introns between them and 120bp intronic flanking sequences before exon 4 and after exon 7 were produced by Synbio Technologies as the plasmid pUC57-dhGao47. pUC57-dhGao47 was used as a template for PCR-amplification of two fragments: one with the primer set LHAdGao47fw / LHAdGao47rev and the other with the primer set RHAdGao47fw / RHAdGao47rev. The 650bp and 430bp PCR products amplified respectively were mixed with pHD-ScarlessDsRed pre-digested with SapI and AarI restriction enzymes and circulated using the NEBuilder HiFi DNA Assembly Cloning Kit. The resultant donor plasmid pLdhGao47R contains humanized exons 4–7 of *Gαo* flanked with short 130bp LHA and RHA and the piggyBac transposon marked with 3xP3-DsRed inserted into the duplicated TTAA sequence in exon 6.

#### Plasmids providing expression of gRNAs under the control of the Drosophila U6:3 promoter

CRISPR targets sites were identified using Target Finder (Gratz et al., 2014), targetfinder.flycrispr.neuro.brown.edu/.

Complimentary oligonucleotides GTCGGACTTTAAACAATATCGAC and AAACGTCGATATTGTTTAAAGTC, GTCGGCCAGCAGCTCCTCCGAGA and AAACTCTCGGAGGAGCTGCTGGC, GTCGTGGCAGGACGCCGGTGTCC and AAACGGACACCGGCGTCCTGCCA, GTC**G**GCAAACAACCTGCGCGGCTG and AAACCAGCCGCGCAGGTTGTTTGC, GTC**G**GACCACTCACCTGTGTATT and AAACAATACACAGGTGAGTGGTC, GTC**G**TTTCCTGGACGATTTGGAT and AAACATCCAAATCGTCCAGGAAA were annealed and cloned into pCFD3-dU6:3gRNA (Addgene, #49410) which was digested with BbsI. 6 plasmids from pCFD-gRNA1 to pCFD-gRNA6 were constructed. Set plasmids pCFD-gRNA1, pCFD-gRNA2, pCFD-gRNA3 combined with the donor plasmid pLdhGao23R were used for the first round of transgenesis; pCFD-gRNA4, pCFD-gRNA5, pCFD-gRNA6 together with the donor plasmid hdGao47ScarlessDsRed – for the second one.

### Flies, germline transformation

Flies were maintained at 25°C on the standard medium. The strain *y[1] sc[*] v[1] sev[21]; P{y[+t7*.*7] v[+t1*.*8]=nos-Cas9*.*R}attP2* expressing Cas9 in the germline under the control of the *nos* promoter (Bloomington Drosophila Stock Center (BDSC), stock no. 78782) was used for germline transformation in the first round of transgenesis. The resultant fly stock *Gαo[h23ex4aa-1]* was combined with *P{y[+t7*.*7] v[+t1*.*8]=nos-Cas9*.*R}attP2* and used for the second round of transgenesis. The strain *w[1118]; In(2LR)Gla, wg[Gla-1]/CyO; Herm{3xP3-ECFP,alphatub-piggyBacK10}M10* expressing PiggyBac transposase (BDSC, stock no. 32073) was used for excision of the PiggyBac-based marker. The resultant alleles and their derivatives were balanced over *CyO*.

Germline transformation was performed as described in the Gompel’s lab protocol (gompel.org/wp-content/uploads/2015/12/Drosophila-transformation-with-chorion.pdf). Embryos were injected with the donor plasmid (500 ng/μl) and three gRNA plasmids (100 ng/μl each).

Transformants were selected under a fluorescence stereomicroscope (Zeiss SteReEO Discovery.V8) using the Filter Set 43 HE for DsRed fluorescent dye detection (excitation BP 550/25; emission BP 605/70).

### Molecular analysis

Genomic DNA was isolated from individual flies of different genotypes as described previously (Gloor et al., 1993). PCR analysis was carried out with different primer sets (Supplementary Table 1, Fig. 2B) using Phusion High-Fidelity DNA Polymerase following the manufacturer’s instructions (New England Biolabs).

Total RNA was isolated with the NucleoSpin RNA kit (Macherey-Nagel) from 30 adult flies for each sample. cDNA was synthesized by priming with oligo-dT with RevertAid Reverse Transcriptase (Thermo Fisher Scientific) following the manufacturer’s instructions. 958bp PCR products including the region from exon 2 to exon 7 were amplified from cDNA with the primers dGaomRNAfw and dGaomRNArev.

### Immunochemistry and microscopy

The following primary antibodies were used: mouse monoclonal antibody against bovine Gαo (1:20 dilution, Santa Cruz Biotechnology, sc-13532) and rabbit anti-dGao (1:50; (Luchtenborg et al., 2014)). Secondary antibodies were donkey anti-mouse Cy3 and donkey anti-rabbit 488 (Jackson ImmunoResearch) used at the 1:300 dilution.

Ventral nerve cords and brains from third instar larvae were fixed with 4% paraformaldehyde in PBS, permeabilized in 0.5% NP-40 and immunostained in 0.2% Tween 20 in PBS. Coverslips were finally mounted with Vectashield (Vector Labs) for microscopy analysis. Humanized and wild-type (control) larval tissues were stained simultaneously in the same vial in order to control the immunostaining specificity.

Fluorescent images were acquired with a Zeiss LSM 800 Airyscan confocal microscope and the images were reconstructed from Z-stacks using ZEN blue software. All images were processed using the same confocal settings.

## Competing interests statement

The authors declare no competing interests.

## Acknowledgements

The work was supported by the Swiss National Science Foundation grant #31003A_175658 to VLK.

